# Ion Dynamics Underlying the Seizure Delay Effect of Low-Frequency Electrical Stimulation

**DOI:** 10.1101/2025.04.01.646594

**Authors:** Guillaume Girier, Isa Dallmer-Zerbe, Jan Chvojka, Jan Kudláček, Přemysl Jiruška, Jaroslav Hlinka, Helmut Schmidt

## Abstract

The biological mechanisms underlying the spontaneous and recurrent transition to seizures in the epileptic brain are still poorly understood. As a result, seizures remain uncontrolled in a substantial proportion of patients. Brain stimulation is an emerging and promising method to treat various brain disorders, including drug-refractory epilepsy. Selected stimulation protocols previously demonstrated therapeutic efficacy in reducing the seizure rate. The stimulation efficacy critically depends on chosen stimulation parameters, such as the time point, amplitude, and frequency of stimulation. This study aims to explore the neurobiological impact of 1Hz stimulation and provide the mechanistic explanation behind its seizure-delaying effects. We study this effect using a computational model, a modified version of the Epileptor-2 model, in close comparison with such stimulation effects on spontaneous seizures recorded *in vitro* in a high-potassium model of ictogenesis in rat hippocampal slices. In particular, we investigate the mechanisms and dynamics of spontaneous seizure emergence, the seizure-delaying effect of the stimulation, and the optimal stimulation parameters to achieve the maximal anti-seizure effect. We show that the modified Epileptor-2 model replicates key experimental observations, and captures seizure dynamics and the anti-seizure effects of low-frequency electrical stimulation (LFES) observed in hippocampal slices. We identify the critical thresholds in the model for seizure onset and determine the optimal stimulation parameters – timing, amplitude, and duration – that exceed specific thresholds to delay seizures without triggering premature seizures. Our study highlights the central role of sodium-potassium pump dynamics in terminating seizures and mediating the LFES effect.

**Author Summary:** This study investigates the mechanisms by which low-frequency electrical stimulation can suppress epileptic seizures. Epilepsy patients often do not respond to pharmacological treatment, necessitating alternative approaches, such as brain stimulation. Using a combination of computational modeling and in vitro experiments on rat hippocampal slices, we examine how periodic stimulation at 1 Hz influences seizure occurrence. Our results show that carefully timed low-frequency stimulation can delay seizure onset by modulating neuronal excitability, largely through the action of the Na-K-pump that maintains ion homeostasis. We employ a modified version of the Epileptor 2 model to reproduce the protective effects seen experimentally. By systematically varying stimulation parameters, we identify conditions that effectively delay seizures, helping to explain the antagonistic effects of stimulation observed by previous studies. Overall, this work advances our understanding of how low-frequency electrical stimulation interacts with intrinsic neuronal mechanisms to prevent seizures, thus offering a potential target for more effective neuromodulation strategies in drug-resistant epilepsy.

## Introduction

Epilepsy is a common neurological disorder with an incidence rate of 61.4 per 100,000 person-years (1, 2). Around 30% of epilepsy patients do not respond to anti-seizure medications (3). Furthermore, only 4.5% of patients are eligible for brain surgery, which is a second-line treatment of epilepsy and a highly invasive procedure (4). Thus, there is a need for the development of new and safe treatment options for epilepsy (5).

Brain stimulation has proven its effectiveness in reducing the seizure rate (6, 7). However, this effectiveness is highly patient- and stimulation parameter-specific, calling for a personalized approach to brain stimulation and epilepsy treatment in general (5, 7). Moreover, the exact cellular and network effect of brain stimulation remains insufficiently understood. This precludes a rational design of stimulation protocols, which is currently replaced by trial-and-error procedures in both the development of new stimulation protocols, as well as in clinical practice (7, 8). To overcome these current challenges in epilepsy treatment, computational modeling has proven advantageous for gaining mechanistic insight and allowing for a safe, inexpensive, and extensive exploration of treatment parameters in the virtual environment (8–10).

In the context of brain stimulation treatment for epilepsy, Low-Frequency Electrical Stimulation (LFES) has been reported to have an inhibitory effect on epileptic activity in models of acute seizures, chronic epilepsy models (11, 12), and in patients with medically intractable seizures (13–16). It thus could serve as a candidate for a new treatment option in epilepsy. However, it appeared that LFES could also trigger epileptic activity (7, 14, 17, 18). Previous studies have shown how this dual effect of stimulation in epileptic tissue is critically dependent on the timing, duration, and amplitude of stimulation, e.g., such that stimulation early after seizure offset would have a seizure-delaying effect while acting in a seizure-promoting fashion when delivered close to an expected seizure onset (9, 19, 20).

The goal of the current study is to investigate and explain the effects of LFES and predict its optimal parameters to delay seizures in local-field potential recordings (LFP) of hippocampal rat brain slices under high-potassium condition, a commonly used animal model of focal epilepsy. To understand the spontaneous seizure dynamics arising from the high-potassium conditions, a biophysical model was employed that is capable of tracking the potassium dynamics, as well as other ion dynamics, linked to the neuronal population activity of excitatory and inhibitory cell populations as recorded by the LFP signal. While various studies have investigated the effects of high-frequency stimulation in mathematical models of epilepsy (20–23), only a few have specifically addressed the impact of low-frequency electrical stimulation (LFES) (24), particularly within frameworks such as the Epileptor-2 model (12, 25).

First, we focus on the emergence of spontaneous seizure-like events recorded in hippocampal slices, and the repeated transitions between the interictal and ictal state. Concurrently, the model parameters are optimized to reproduce the different characteristics of the individual slices, explaining their heterogeneity. Second, we demonstrate that the model can accurately capture the seizure-delaying effect of LFES. By stabilizing the system into a seizure-free attractor, LFES is shown to stop extracellular potassium accumulation by sustaining the activity of sodium-potassium pumps, thereby preventing seizure generation.

## Materials and Methods

### Experimental Design

Brain slices were obtained from four male Wistar rats weighing approximately 200g. The animals were kept in an enriched environment with 12/12 hours of light and darkness and regularly checked for health status prior to the experiment. The animals were anesthetized (80 mg/kg ketamine, 25 mg/kg xylazine) and decapitated. The brains were extracted from the skull and placed into an ice-cold, oxygenated protective solution – sucrose-based artificial cerebrospinal fluid – which contained (in mM) 189 mM sucrose, 10 glucose, 2.5 KCl, 0.1 CaCl_2_, 5 MgCl_2_, 26 NaHCO_3_, and 1.25 NaH_2_PO_4_*H_2_0.

The brain was then cut into saggital slices of 350 *μ*m thickness using a vibratome (Campden Instruments, Loughborough, UK). The CA3 region was then cut off by a scalpel. Then, the hippocampal slices were moved to a holding chamber at room temperature filled with normal artificial cerebrospinal fluid (ACSF) and continuously aerated with a humidified mixture of 95% O_2_ and 5% CO_2_. The ACSF contained (in mM) 125 NaCl, 26 NaHCO_3_, 3 KCl, 2 CaCl_2_, 1 MgCl_2_, 1.25 NaH_2_PO_4_, and 10 glucose. After at least 60 minutes, a slice was transferred to a recording interface chamber at 34 ± 1°C containing normal ACSF. They were left at rest for 10 minutes before experiments started. Local field potentials were recorded using a glass micropipette (diameter 10 - 15 *μ*m) filled with ACSF and inserted in the stratum pyramidale of the CA1 subregion. The signal was amplified using Model 3000 AC/DC Differential Amplifier (A-M Systems, Inc., Carlsborg, Washington, USA), digitized at 10 kHz sampling frequency using Power1401 (CED, Cambridge, UK) and recorded using Spike2 software (CED, Cambridge, UK). To induce epileptic activity, we added KCl solution at first in large steps (1 - 3 mM) until a total concentration of 6.5 - 7 mM was reached. After that, we gradually increased the concentration (0.2–0.5 mM at a time) until spontaneous seizure-like episodes appeared. The experimental protocol included perturbations of the CA1 network with regularly delivered stimuli to Schaffer collaterals (input to CA1) with a frequency of 1 Hz. The stimuli were electrical pulses of 200 *μ*s duration using a bipolar silver wire electrode and isolated constant current stimulator (Digitimer Ltd., Welwyn, UK). The stimulation was initiated immediately after the end of a seizure-like event, at the beginning of the inter-seizure interval (ISI). A stimulated ISI was typically followed by an ISI without stimulation and vice versa. In the analysis, each stimulated ISI was matched to the preceding control ISI. The stimulation duration ranged from a few seconds to nearly two minutes. Table 1 summarizes basic data characteristics. The signals were subsequently analyzed using custom-written Matlab scripts. The data were recorded previously (19) and reanalyzed for the purposes of the present study. All experimental procedures were approved by the Czech Academy of Sciences Ethics Committee and the Ethics Committees of the Second Faculty of Medicine (Project Licence No. MSMT-31765/2019-4).

**Table 1.** Data characteristics of each slice. K_*bath*_: final potassium concentration per slice, which was sufficient to elicit seizure-like events; n_*seiz*_ baseline: number of spontaneous seizures before the stimulation experiment was started; ISI baseline: inter-seizure interval length (mean ± standard deviation) before the stimulation experiment; n_*seiz*_ stim: number of seizure trials with stimulation (where pre-stim seizure trials without stimulation served as a matched control for assessing for relative ISI changes); ISI stim: ISI with stimulation, ISI pre-stim: ISI without stimulation preceding an ISI with stimulation, used as control.

| | $K_{bath}$ | $n_{seiz}$ baseline | ISI baseline | $n_{seiz}$ stim | ISI stim | ISI pre-stim | stim length |
| --- | --- | --- | --- | --- | --- | --- | --- |
| slice 1 | 9.5 mM/L | 5 | 64 ± 3 s | 28 | 78 ± 26 s | 56 ± 5 s | 51 ± 24 s |
| slice 2 | 8.6 mM/L | 8 | 58 ± 12 s | 15 | 73 ± 23 s | 46 ± 7 s | 40 ± 22 s |
| slice 3 | 8 mM/L | 10 | 56 ± 2 s | 21 | 75 ± 22 s | 53 ± 6 s | 58 ± 25 s |
| slice 4 | 5.3 mM/L | 6 | 43 ± 9 s | 22 | 45 ± 10 s | 34 ± 6 s | 27 ± 12 s |

### Statistical Analysis

To demonstrate the seizure-delaying effect due to LFES in the experimental data, we computed the Spearman correlation between the length of stimulation and the seizure delay for each slice. Seizure delay was calculated as percentage change of the stimulated ISI with respect to the control ISI (which was the unstimulated ISI immediately preceding each respective stimulated ISI, i.e. “pre-stim ISI” in Table 2). We report the correlation coefficient *r* and the associated p-values. The model analysis did not require any statistical analysis.

**Table 2.**
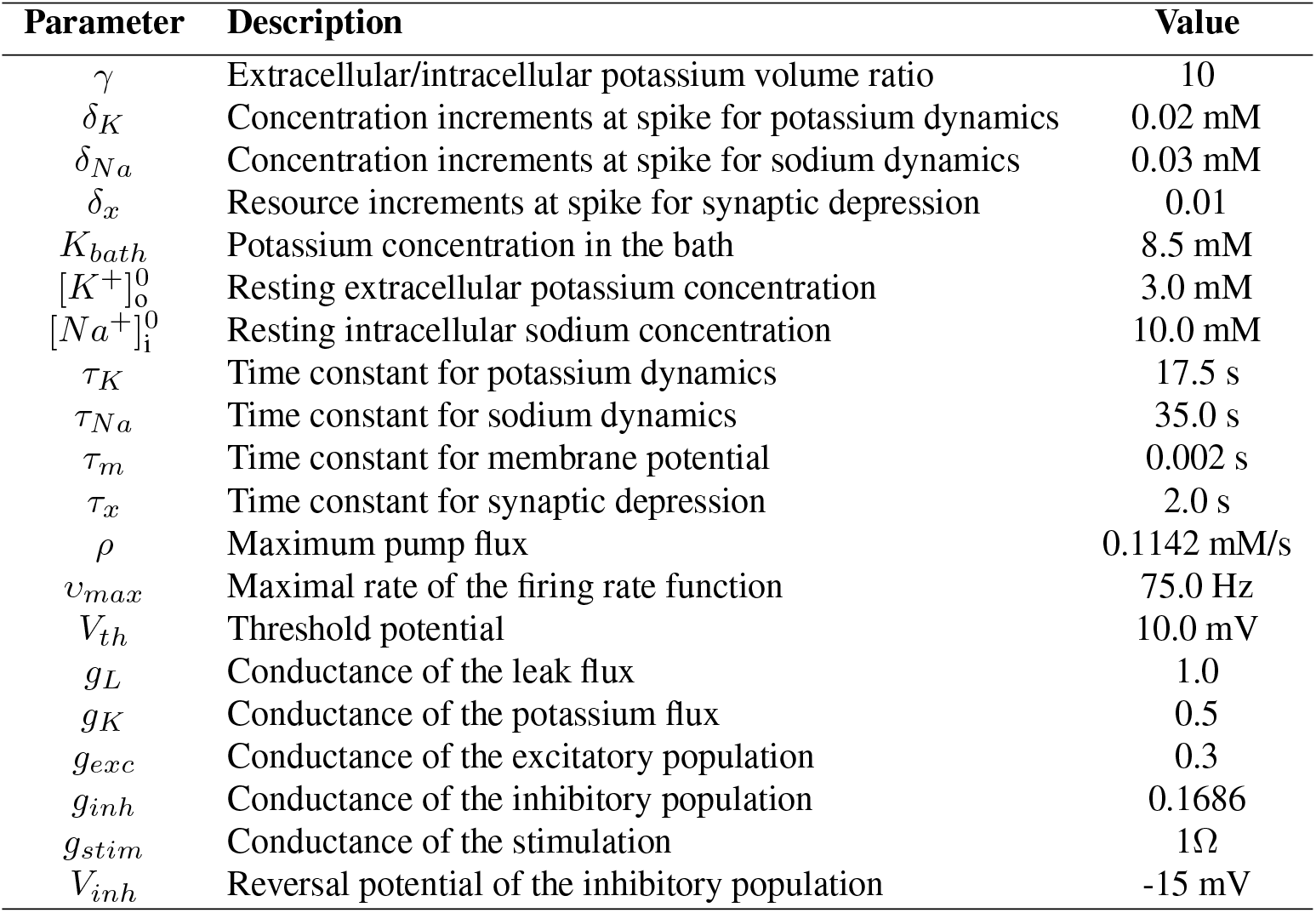
Fixed parameter set used in Section Results.

| Parameter | Description | Value |
| --- | --- | --- |
| $\gamma$ | Extracellular/intracellular potassium volume ratio | 10 |
| $\delta_K$ | Concentration increments at spike for potassium dynamics | 0.02 mM |
| $\delta_{Na}$ | Concentration increments at spike for sodium dynamics | 0.03 mM |
| $\delta_x$ | Resource increments at spike for synaptic depression | 0.01 |
| $K_{bath}$ | Potassium concentration in the bath | 8.5 mM |
| $[K^+]_o^0$ | Resting extracellular potassium concentration | 3.0 mM |
| $[Na^+]_i^0$ | Resting intracellular sodium concentration | 10.0 mM |
| $\tau_K$ | Time constant for potassium dynamics | 17.5 s |
| $\tau_{Na}$ | Time constant for sodium dynamics | 35.0 s |
| $\tau_m$ | Time constant for membrane potential | 0.002 s |
| $\tau_x$ | Time constant for synaptic depression | 2.0 s |
| $\rho$ | Maximum pump flux | 0.1142 mM/s |
| $v_{max}$ | Maximal rate of the firing rate function | 75.0 Hz |
| $V_{th}$ | Threshold potential | 10.0 mV |
| $g_L$ | Conductance of the leak flux | 1.0 |
| $g_K$ | Conductance of the potassium flux | 0.5 |
| $g_{exc}$ | Conductance of the excitatory population | 0.3 |
| $g_{inh}$ | Conductance of the inhibitory population | 0.1686 |
| $g_{stim}$ | Conductance of the stimulation | 1Ω |
| $V_{inh}$ | Reversal potential of the inhibitory population | -15 mV |

The ISI was defined as the time interval in seconds between the end of one seizure and the beginning of the next. In the data, seizure offsets were automatically identified by finding the minimum in the LFP (using Matlab R2021a’s *islocalmin()* function with ‘MinProminence’ set to half the signal span and a ‘MinSeparation’ 20 seconds, applied to the smoothed signal; *smooth()* with window size of one second). Seizure onsets were identified by finding the steep increase of LFP (using *findchangepts()* function with ‘Statistic’ set to ‘linear’) in the time window of up to 30 seconds before each identified seizure offset.

### Model Design

In this study, we aimed to investigate a minimal biophysical model of a neuronal population that: 1) exhibits epileptiform activity similar to our data, 2) incorporates potassium concentration dynamics, and 3) can produce seizure delay in response to low-frequency electrical stimulation.

We chose a modified version of the Epileptor-2 model (25):

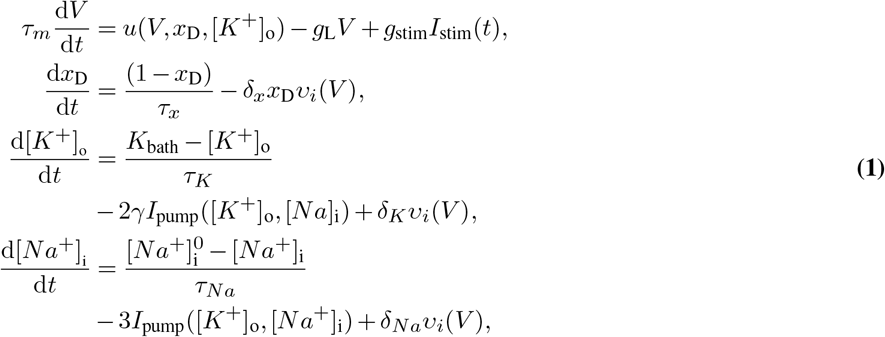

where *V* is the average membrane potential of pyramidal cells, *x*_D_ is the synaptic resource, and [*K*^+^]_o_ and [*Na*^+^]_i_ are the extracellular potassium and intracellular sodium concentrations, respectively.

The membrane potential *V* is driven by the potassium concentration [*K*^+^]_o_ and synaptic input, which is concisely expressed by

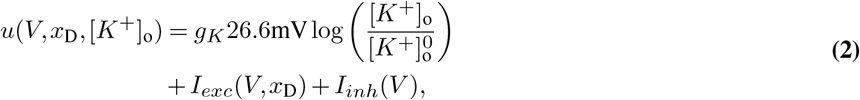

where the first term is the potassium depolarizing current, and *I*_*exc*_ and *I*_*inh*_ are the excitatory and inhibitory synaptic inputs to the excitatory population. The excitatory synaptic input *I*_*exc*_ is defined as:

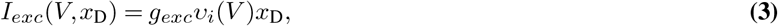

with *g*_*exc*_ being the conductance of excitatory synapses, and *υ*_*i*_(*V*) being the firing rate of the excitatory population. We note here that the excitatory synaptic input is current-based, since the reversal potential for excitatory synapses is so large that its difference from the membrane potential can be considered roughly constant (26). The inhibitory synaptic input *I*_*inh*_ is defined as (27):

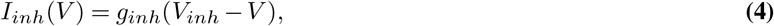

with *g*_*inh*_ being the conductance of inhibitory synapses, and *V*_*inh*_ their reversal potential.

The firing rate function *υ*_*i*_(*V*) is chosen to be a sigmoid function with threshold *V*_*th*_ and maximum firing rate *υ*_*max*_:

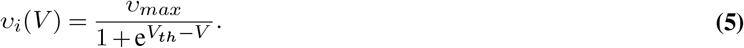

The excitatory population is also subjected to low-frequency stimulation, *I*_*stim*_(*t*). Each stimulus is represented by a Dirac delta function, and the sequence of stimuli thus reads *I*_*stim*_(*t*) = *A*Σ_*n*_ *δ*(*t* − *t*_*n*_), where *t*_*n*_ represents stimulation times (once per second in our protocol), and *A* is the stimulation amplitude. The parameter *g*_*stim*_ was added to ensure consistency of units. Lastly, the activity of the Na-K pump is computed as follows (28):

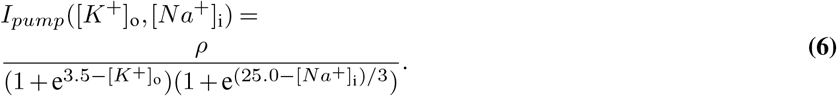

We made two modifications to the original Epileptor-2 model (25), because the original Epileptor-2 was not sufficient to reproduce the seizure delay effect. First, we changed the firing rate function to a simple sigmoid function, which allows for non-zero firing rates even at small values of the membrane potential. The second modification concerns the implementation of inhibition. The original Epileptor-2 model features a non-depressing inhibitory neuronal population whose activity is proportional to that of the excitatory neuronal population, thus providing a tight balance between the inhibitory and excitatory population. Here, however, we chose an inhibitory population that exhibits a constant firing rate and acts on the excitatory population via conductance-based synapses. Since the reversal potential *V*_*inh*_ is close to the resting potential at low [*K*^+^]_*o*_ concentrations, inhibition doesn’t produce strong hyperpolarizing synaptic potentials. Rather, it has a shunting effect that is roughly proportional to the depolarization of the membrane. Experimental and computational evidence suggests that shunting inhibition is prevalent in the hippocampus, and that it plays a crucial role in the robustness of *γ*-oscillations (29, 30).

One of the objectives is to investigate the effect of potassium on the neuronal population. In the full model, the sodium and potassium concentrations are slowly changing variables that can be assumed to be roughly constant on the much faster time scales that govern the dynamics of *V* and *x*_D_. Therefore, to describe the characteristic dynamics of *V* and *x*_D_, we will assume that [*K*^+^]_*o*_ and [*Na*^+^]_*i*_ are time-invariant, leading to the following set of equations:

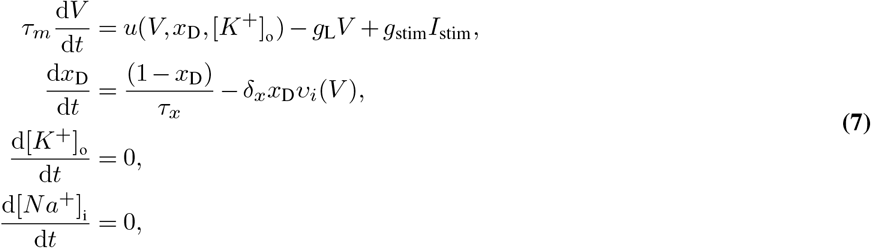

This system allows us to obtain the bifurcation structure of the system with respect to the ion concentration which can be studied as parameters.

In this analysis, we consider two scenarios: 1) using parameter sets fitted to the experimental data (for slices 1 to 4) and 2) using an arbitrary parameter set to explore LFES in greater detail, referred to as slice 0. The parameters that are kept constant throughout the analysis are shown in Table 2.

The parameters specific to the slice 0 and the ones specific to the experimental slices 1-4 are shown in Table 3.

**Table 3.**
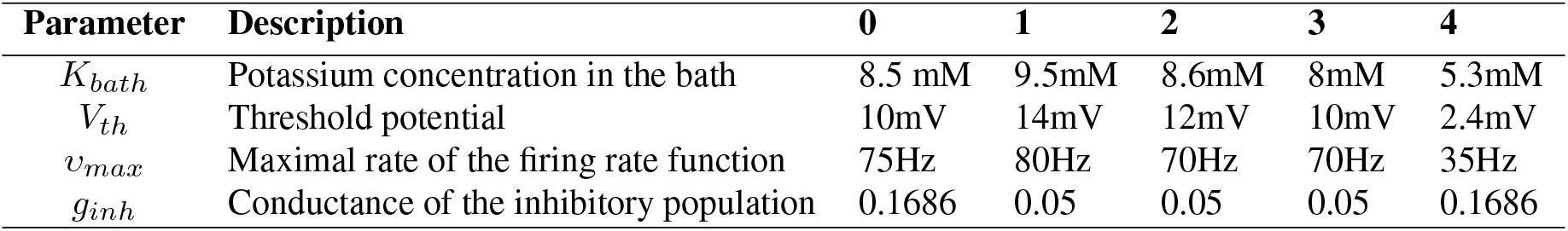
Specific parameter set used to fit the experimental data.

| Parameter | Description | 0 | 1 | 2 | 3 | 4 |
| --- | --- | --- | --- | --- | --- | --- |
| $K_{bath}$ | Potassium concentration in the bath | 8.5 mM | 9.5mM | 8.6mM | 8mM | 5.3mM |
| $V_{th}$ | Threshold potential | 10mV | 14mV | 12mV | 10mV | 2.4mV |
| $v_{max}$ | Maximal rate of the firing rate function | 75Hz | 80Hz | 70Hz | 70Hz | 35Hz |
| $g_{inh}$ | Conductance of the inhibitory population | 0.1686 | 0.05 | 0.05 | 0.05 | 0.1686 |

### Code and Data Accessibility

All code supporting the findings of this study are publicly available at https://github.com/GuillaumeGirier/Low-Frequency-Electrical-Stimulation-and-Seizure-Delay. The experimental data will be made available at the time of publication.

## Results

### The emergence of spontaneous seizure transitions

We first examine the spontaneous transitions into seizures by analyzing how the model parameters influence the duration of seizures and ISIs. Fig. 1.A shows four example LFP recordings, obtained from the experimental setup described in the Methods section. It shows the heterogeneity in terms of ISI and seizure duration between slices, which is dependent on the amount of added potassium. Based on this observation, we decided to match the recordings with the model in terms of seizure duration and ISI.

**Fig. 1.**
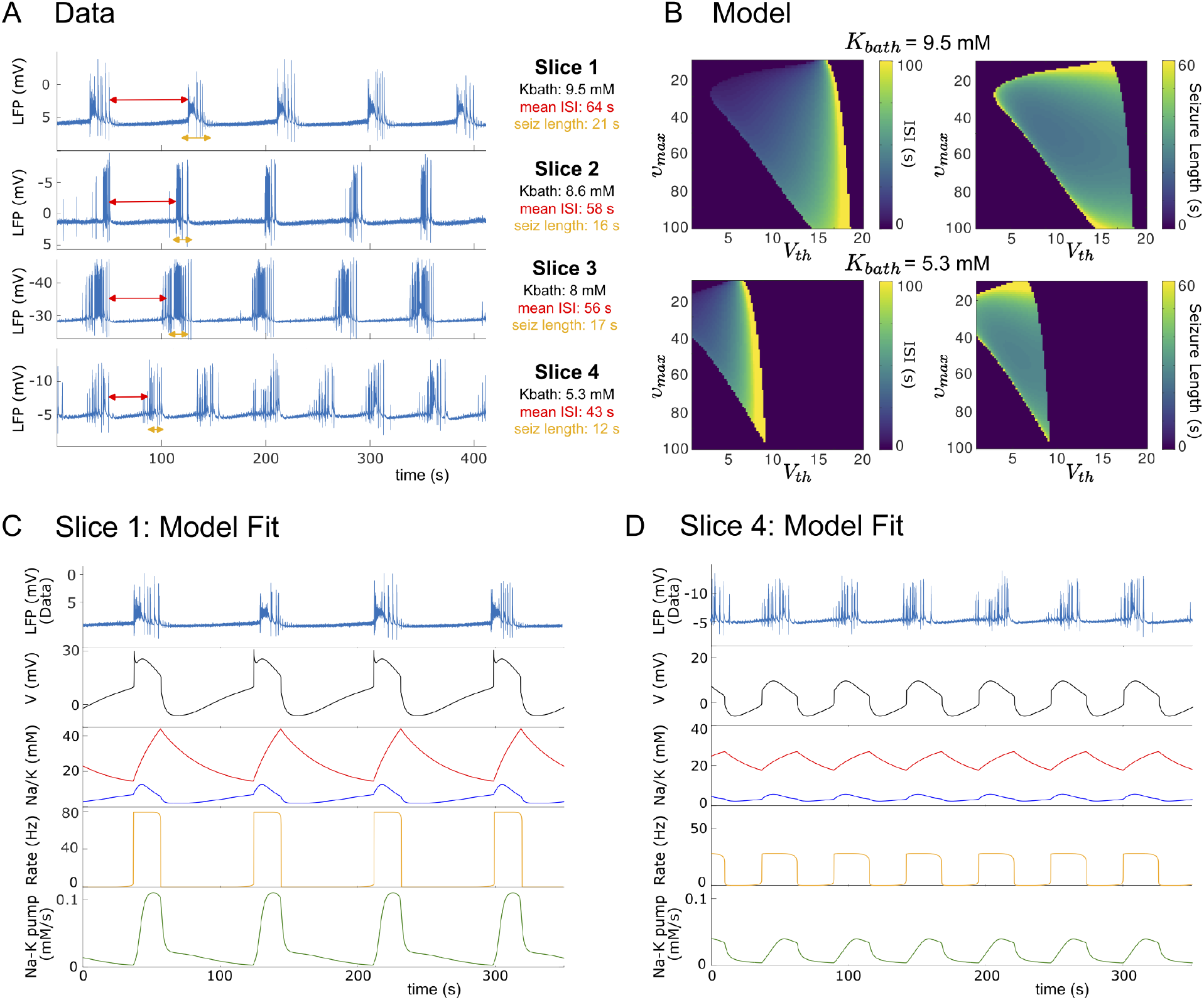
Fitting model ISI and seizure duration to the data. (A) Time series of data from different slices. Red double arrows indicate ISIs, and orange double arrows indicate seizures. (B) Heatmaps of ISI and seizure duration with respect to the parameters *υ*_*max*_ and V_*th*_, for *K*_*bath*_ = 9.5 and *K*_*bath*_ = 5.3. (C) Model fit to slice 1. (D) Model fit to slice 4. In these panels, red curves refer to [*Na*^+^]_*i*_ concentration, and the blue curves refer to [*K*^+^]_*o*_ concentrati
on. For the specific parameters for each slice, see Table 3.

The Epileptor-2 model contains various parameters that can be modified to change the temporal structure of the epileptic activity and thus replicate the experimental observations. In our case, we focus on the firing threshold *V*_*th*_ and the maximum firing rate *υ*_*max*_. It is reasonable to assume that slices differ in specific characteristics (31–34), such as cellular excitability and network connectivity, due to intrinsic biological heterogeneity and variations in preparation protocols. This variability underscores the importance of systematically analyzing the parameters *V*_*th*_ and *υ*_*max*_ to account for the diversity observed in the dataset.

The heatmaps in Fig. 1.B determine the length of the ISI and seizures as *V*_*th*_ and *υ*_*max*_ are varied, demonstrated for the *K*_*bath*_ values of slices 1 and 4. We note here that if the ISI and seizure length are equal to zero, the model exhibits constant activity with either a low or high firing rate. For fixed values of *υ*_*max*_, the length of the ISI increases and the seizure length decreases as *V*_*th*_ increases.

Using the heatmaps in Fig. 1.B, we calibrate the model to match two examples from Fig. 1.A (slice 1 and 4). In Fig. 1.C and D, we first show a time series of the LFP from the data, then a time series of the membrane potential from the model. One can notice the similarity in structure of the epileptic activity of the model: a slow depolarization that characterizes the ISI, then a fast depolarization that starts the seizure. The seizure itself is characterized by initially further depolarization, and subsequent repolarization until the membrane potential reaches its minimum, which terminates the seizure. It exhibits very similar sodium and potassium concentrations as determined experimentally (35). We note here that the resting sodium concentration in the model is 10mM, which is representative of the normal physiological state. In addition to characterizing the dynamics of potassium and sodium ions, the model provides access to the firing rate and the activity of the Na-K pumps. Shortly before a seizure, the firing rate shows a modest increase, followed by a rapid upturn at the beginning of the seizure. At the end of the seizure, the firing rate declines abruptly. In parallel, Na-K pump activity gradually increases during the seizure. At the seizure end, pump activity initially decreases rapidly as the membrane potential returns to baseline. After reaching baseline, the activity continues to decline but at a slower steady rate.

In Fig. 2.A, we first test different values of *K*_*bath*_ to study their effect on the time series. The potassium concentration in the surrounding bath, *K*_*bath*_, is increased instantaneously at the vertical dashed line. We note that the addition of potassium allows the appearance of seizures only when a specific concentration threshold is exceeded, then the ISI progressively reduces and the seizure increases in length until the added potassium stabilizes the potential at a high frequency value. This status could correspond to so-called *status epilepticus*, i.e. an epileptic seizure prolonged for more than 5 minutes or by repeated seizures without regaining consciousness between them (36–38). We remark here that the alternative scenario of depolarisation block is not captured by our model, because the firing rate saturates at high values of the membrane potential, which represents intense firing activity. Note that the length of ISI and seizure duration depend also on the *V*_*th*_ and *υ*_*max*_, which is fitted individually to each slice in Fig. 1.C,D. For each setting of *V*_*th*_ and *υ*_*max*_ the effect of increasing *K*_*bath*_ corresponds to the one shown in Fig. 2.A. Comparing *K*_*bath*_ concentrations between slices 1 and 4, we found longer ISI and shorter seizures for higher *K*_*bath*_, while increasing *K*_*bath*_ within one slice will lead to shorter ISI and longer seizure length. This discrepancy is resolved when considering that each slice has different characteristics, which corresponds to different parameter combinations placing it in a different point in Fig. 1.B.

**Fig. 2.**
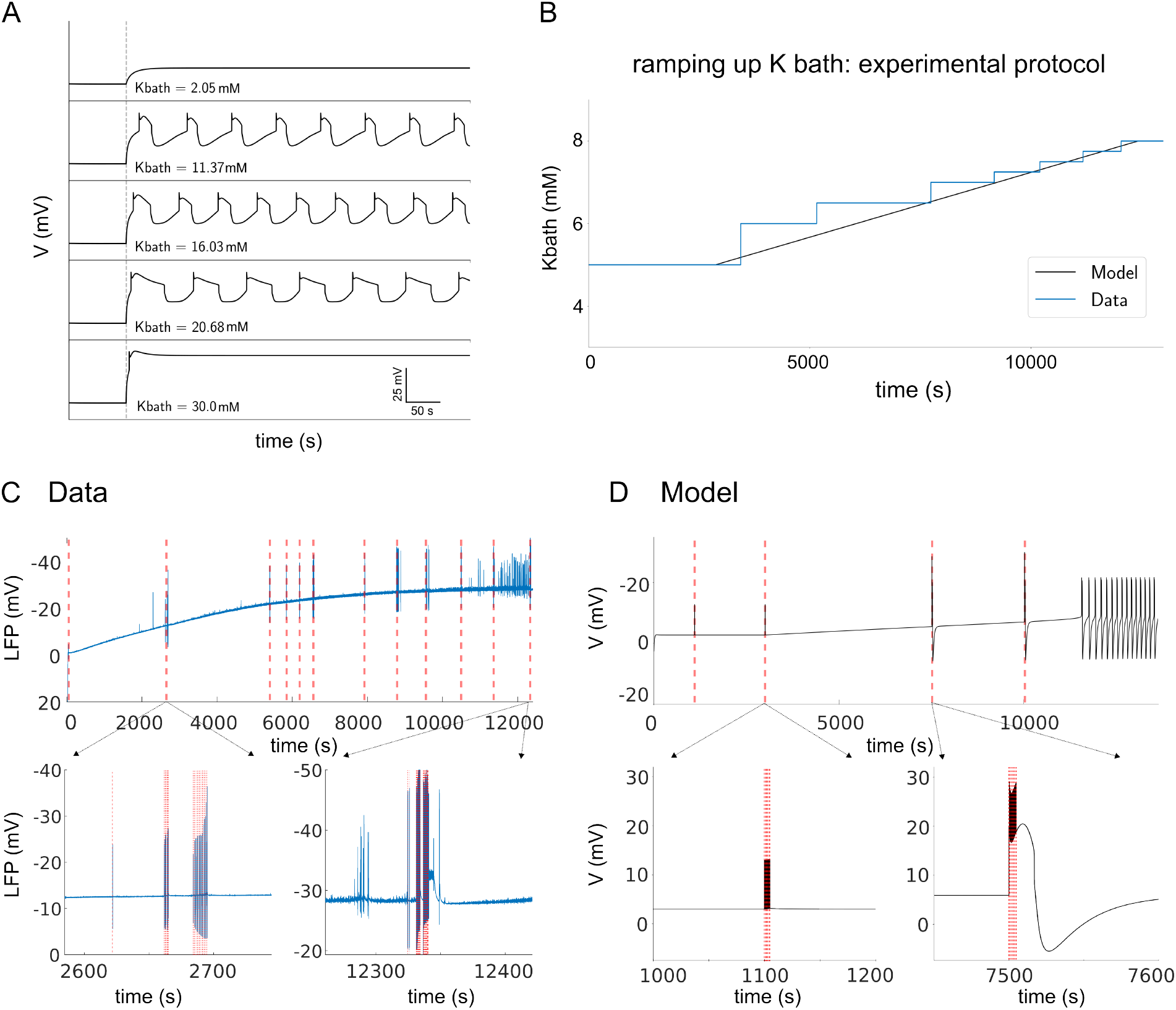
Highlighting the effect of *K*_*bath*_ on the modified Epileptor-2 model compared to the data. (A) Time series of the model for different values of *K*_*bath*_. (B) Superposition of the experimental protocol of sequential addition of potassium performed in the medium containing the slices, with the simulated slow addition protocol. The red vertical dashed lines correspond to the moment when the model is stimulated with a single spike of *A* = 15mA. (C) Effect of the same experimental protocol: the blue time series corresponds to the membrane potential, and the red one to the moment when the experimentalists stimulated the slice. (D) Effect of the slow addition protocol on the model. For panel (C) and (D), zooms on the time series at the moment of the stimulation effect are provided. For the specific parameters, see slice 0 in Table 3.

In the next step, we focus on the potassium addition protocol: in Fig. 2.B, the experimental protocol (blue curve), which corresponds to regular additions in the medium with a delay long enough to allow the potassium to diffuse into the tissue, is superimposed onto our protocol applied to the model (black curve). In the experimental protocol, potassium is added until the first appearance of spontaneous, recurring seizures. However, sometimes the experimenter probed the tissue excitability by stimulating the tissue in order to induce the appearance of seizures, as commonly done in epilepsy research and clinical practice (39). In order to reproduce the experimental setup, we added a constant slow input of potassium to our model. The continuous addition of potassium in the model, as opposed to the step-wise manner during the experiment, is justified by the slow diffusion process at each step, so we can assume that our process is similar. Furthermore, we tested the induction of seizures by external stimulation with spike trains of 30 mA, which are indicated in Fig. 2.D by red dashed vertical lines.

We first examine the effect of stimulations in the experimental protocol. Fig. 2.C shows the recorded LFP from slice 3, overlaid with the different stimulations applied to induce seizures. At low potassium concentrations, the stimulation produced only transient depolarization-repolarization responses, regardless of the number of pulses that were applied. However, at higher potassium concentrations later during the experiments, the stimulation was capable of provoking seizure events if the number of pulses was sufficiently large.

In Fig. 2.D, the experimental protocol is reproduced in the model, showing the membrane potential of the neuronal population. In these simulations, the stimulation consists of a train of spikes delivered at 1 Hz. We observe that when the extracellular potassium concentration is low, the stimulation only produces artifacts without triggering any pathological activity. However, when the potassium concentration is elevated, it is the first spike of the stimulation train that is sufficient to initiate a seizure.

### The anti-seizure effect of LFES

To demonstrate the seizure-delaying effect of LFES, we analyzed the LFP recordings with regards to the difference in length between stimulated ISIs and control ISIs (which are ISIs without stimulation preceding the stimulated ISI, i.e. “pre-stim ISI” Table 2). The seizure delay was calculated as percentage change of the stimulated ISI with respect to the control ISI.

We find that stimulated ISIs were significantly longer than the control ISIs, in each of the slices, as assessed via Spearman correlation (see Fig. 3.B and Table 2; see also Supplemental Material S1). This indicates that the 1 Hz stimulation had indeed a seizure delaying, or anti-seizure, effect. Fig. 3.A shows two examples of control and subsequent stimulated ISI (their beginnings and ends marked by red and green vertical lines, respectively). The longer stimulation in example 2 leads to a substantially longer stimulated ISI as compared to control ISI.

**Fig. 3.**
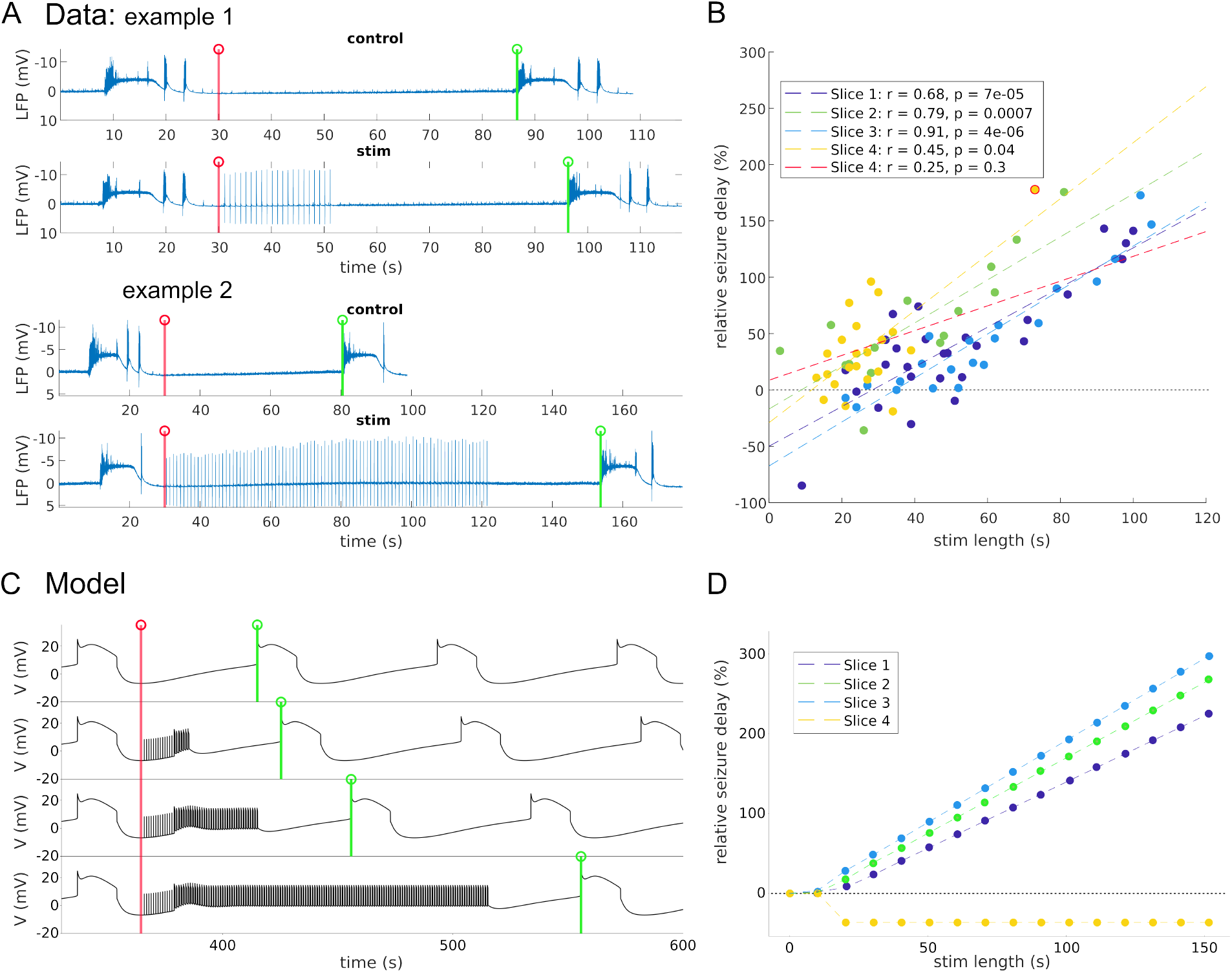
LFES-induced delay phenomena observed from the data and from the model. (A) Time series from experimental data. The beginning (end) of the ISI is marked by a red (green) vertical line. (B) Seizure delay correlation with stimulation length from experimental data, as assessed via Spearman correlation. Seizure delay is measured as the percent change of stimulated ISI relative to control ISI. Each dashed line is a linear regression fit. For slice 4, two scenarios are proposed: one with the whole data (yellow dashed line), and one without outlier (red dashed line). The outlier is circled in red. (C) Time series of the simulated model with standard parameters (“slice 0”) for different stimulation lengths. (D) Seizure delay correlation with stimulation length for the model fitted to slices 1 to 4. For the specific parameters for each slice, see Table 3.

In Fig. 3.C and D, we apply the LFES protocol to the model. In all the simulations, the stimulation is applied right after the seizure. In Fig. 3.C, we can observe that the stimulation pulse does not have an immediate effect for approximately the first 10 to 15s. Only after this initial period, we observe a tangible effect of the stimulation, which is a brief increase in the depolarization rate, and subsequently the convergence of the system onto a new forced equilibrium until the stimulation ends. Thus there exists an optimal time window in which the stimulation is effective, which we investigate further below.

Fig. 3.D shows the effect of stimulation on the ISI in the same way as panel B. For values up to *t*_*stim*_ = 10s, the stimulation does not have an effect on the ISI, in line with the (descriptive) experimental observations in Fig. 3.B that for short stimulation lengths LFES is ineffective. For stimulation periods greater than *t*_*stim*_ = 10s, in the case of slices 1 to 3, the ISI increases linearly with the stimulation period. The model reproduces the same order of slope as observed in the experimental data (Data: Slope_slice 1_ =0.830, Slope_slice 2_ =0.933, Slope_slice 3_ =0.937; Model: Slope_slice 1_ =1.027, Slope_slice 2_ =1.183, Slope_slice 3_ =1.268). This demonstrates its ability to predict the individual effectiveness of the anti-seizure protocol. However, in the case of slice 4, the ISI decreases and then stabilizes, as LFES induces an early seizure. This phenomenon arises from the choice of parameters optimized to fit the experimental data. The impact of LFES on seizure delay as a function of *V*_*th*_ and *υ*_*max*_ will be examined in the Supplemental Material S2.

### Model predictions regarding optimal parameters and the mechanism of the anti-seizure effect

In order to understand the mechanism behind the anti-seizure effect, we first investigate the optimal stimulation parameters — timing, and amplitude — that maximize it. If the stimulation causes a new seizure, then this is considered the end of the ISI, even as the stimulation continues.

We portray the changes to the ISI in a heatmap in Fig. 4. In panel A, we plot the effect of amplitude and onset time while keeping the stimulation duration fixed at 50 s. When the amplitude is low, e.g., *A* = 0.5 mA, the ISI does not change significantly. When the amplitude increases from 2.5 mA to 12.5 mA, the stimulation reduces the ISI. However, when the amplitude exceeds 15 mA and the onset time is 16s or less, the stimulation increases the ISI by 40% to 60% compared to the unperturbed ISI. If the stimulation onset occurs too late (17s or more), then the stimulation creates a new seizure immediately. We can speak here of a critical point for the stimulation amplitude and stimulation onset. In Fig. 4.B, we plot some corresponding time series that are around this critical point. In panels B.1, 2 and 4, the LFES causes premature seizures. In Fig. 4.B.3, stimulation effectively delays the next seizure and pushes the system onto a new forced equilibrium during stimulation. One can also observe that in panel B.4, the LFES does indeed induce a premature seizure, however unlike panels B.1 and 2 where the stimulation amplitude is not high enough to delay subsequent seizures, what follows the first premature seizure is the beginning of a delay process. This implies that even if the stimulation starts too late and triggers an earlier seizure it can delay the next seizure if the amplitude is high enough.

**Fig. 4.**
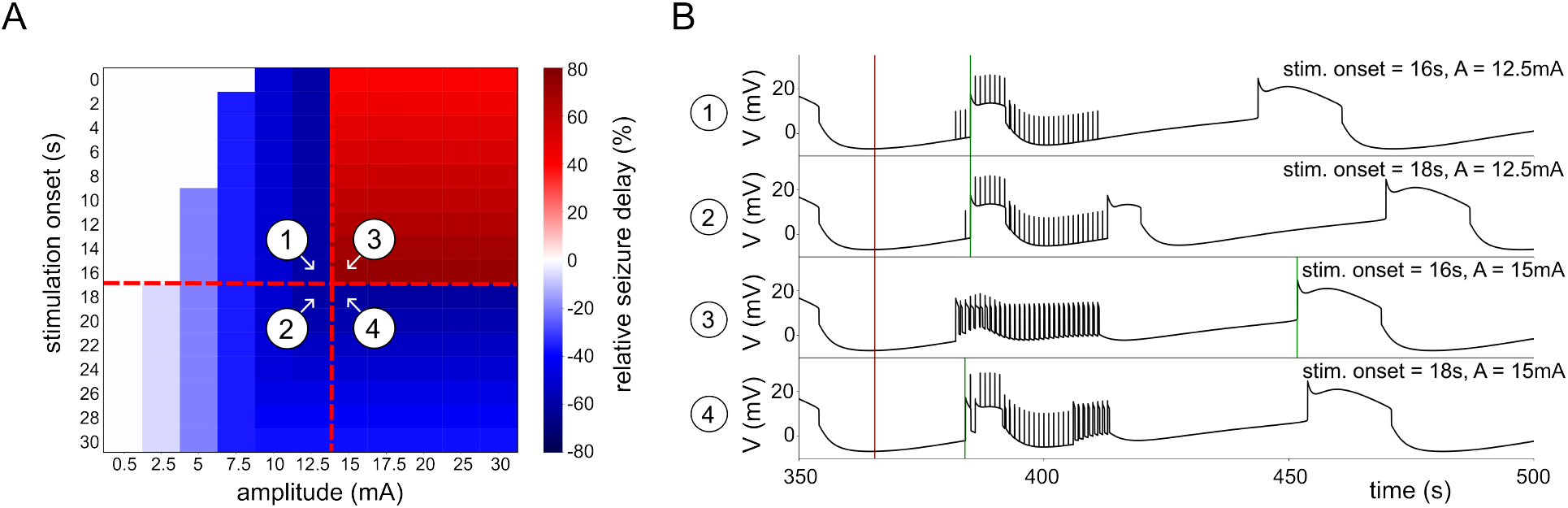
Relative seizure delay as a function of two variables: the amplitude of the signal (mA), and the stimulation onset after the end of the first seizure (s). (A) Relative seizure delay plotted against amplitude and stimulation onset, with the stimulation length fixed at 25s. (B) Time series numbered from 1 to 4 illustrate model behavior around the critical point in (A). For the specific parameters, see slice 0 in Table 3.

Note that our experimental dataset was not sufficient to validate the model predictions regarding optimal parameters. Stimulation amplitude information was not available. Moreover, the stimulation onset time with regard to last seizure offset time varied only between −4.8 and 7.3 seconds with a median of 1.2 seconds (see Supplemental Material S3).

Having identified the critical thresholds for each stimulation parameter, we next aim to elucidate the biological processes underlying the effects of LFES. To achieve this, we analyze the model time series by mapping them onto the Na-K phase plane. To interpret this representation, we highlight the stimulation threshold and employ a bifurcation diagram to delineate the distinct dynamical regimes—namely, ictal and interictal states. This approach allows us to assess whether LFES effectively drives the system away from the ictal into the interictal region.

Fig. 5.A shows one time series obtained from the model, namely the membrane potential, Na/K concentrations, firing rate, and Na-K pump activity, for the case where the LFES is active for 150 s. The stimulation sends the system into a regime where the Na-K pump is active (Na: 25.4 − 25.8 mM, K: 3.4 − 3.7 mM). A stimulation pulse causes an instantaneous increase of both Na and K concentration. Then, the active Na-K pump slowly lowers both again. This requires sufficiently strong Na-K pump activity, otherwise the mechanism is likely to fail.

**Fig. 5.**
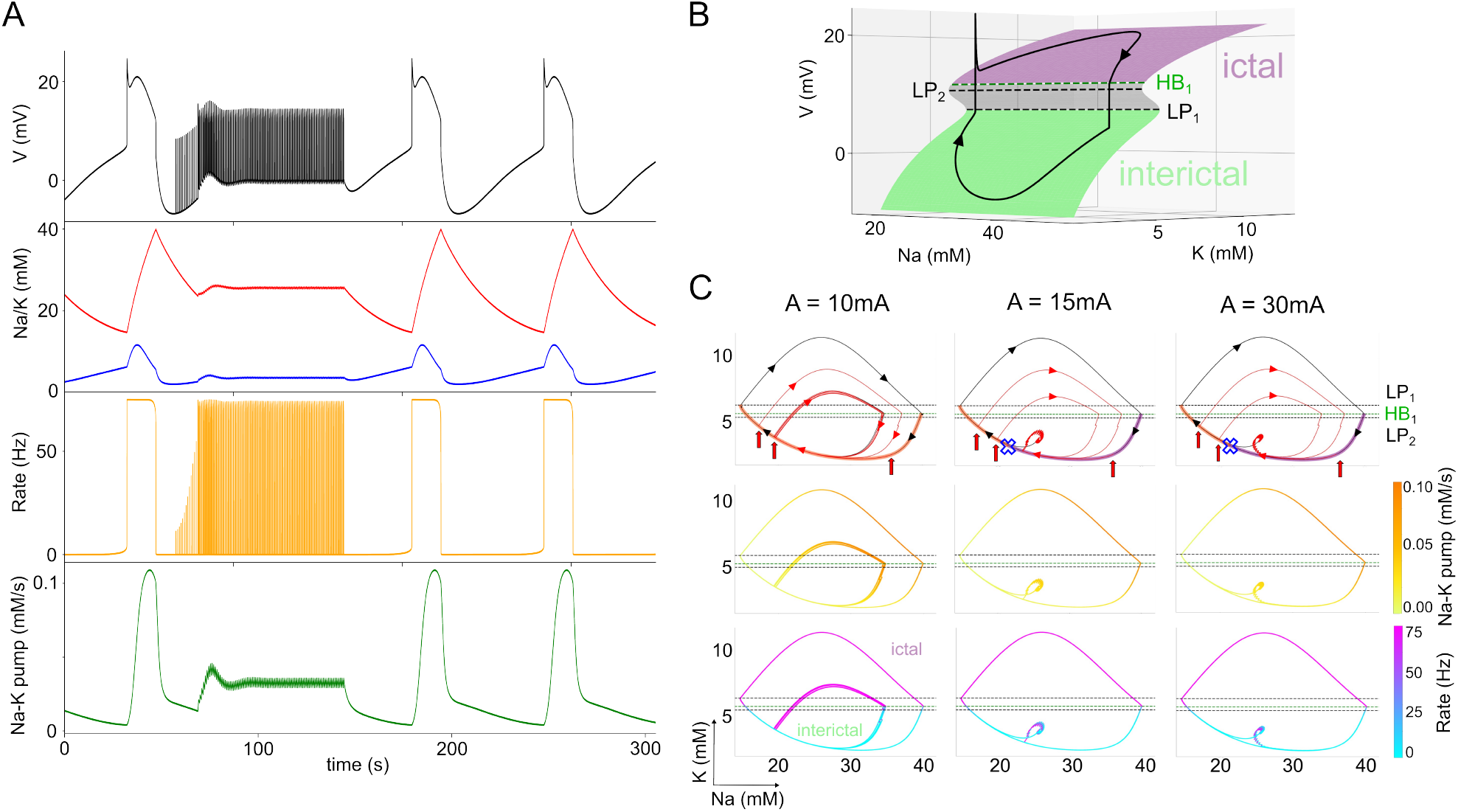
Deciphering the biological mechanisms underlying LFES. (A) Time series demonstrating seizure delay in response to LFES. (B) 3D {[*K*^+^]_*o*_, [*Na*^+^]_*i*_, *V*} bifurcation diagram obtained from the fast-subsystem (Eq. Eq. (7)), overlaid with a non-stimulated solution obtained from Eq. Eq. (1) (black arrowed curve). For the superposition to be in perfect adequacy, we impose *ε* = 0.003 parameter in front of the equation of [*K*^+^] and [*Na*^+^], in such a way, they are slower and follow the bifurcation structure. The stable states are the green and purple surface, which refer respectively to the inter-ictal region and the ictal region. The grey surface represents the unstable states. HB denotes Hopf bifurcation, and LP, fold bifurcation point. (C) Effect of our protocol on Eq. Eq. (1) represented on a {[*Na*]^+^, [*K*]^+^} phase plane respectively for *A* = 10, 15, and 30 mA by a stimulated trajectories (arrowed curves), and colormaps of the activity of Na-K pumps, and the firing rate. The start of each stimulation is represented by a vertical red arrow, and last 150s. Trajectories with LFES are plotted in red, trajectories without LFES are plotted in black. The trajectories highlighted in orange or purple indicates where LFES respectively induces an earlier seizure or delays its onset. The blue cross represents the critical point up to which a stimulation causes a delay, as obtained in Fig. 4. For the specific parameters, see slice 0 in Table 3.

This explains why stimulation that occurs too soon after the previous seizure is ineffective in delaying or suppressing the next seizure: the K concentration is still too low and the Na-K pump is inactive. Only when the K concentration is sufficiently high (above ≈ 2.5 mM), the mechanism starts working.

We now characterize the effects produced by LFES in greater detail. We first focus on the {[*K*^+^]_*o*_, [*Na*^+^]_*i*_, *V*} bifurcation diagram obtained from Eq. (7) (S-shaped, colored surface in Fig. 5.B), which is superimposed with the trajectory of a solution of the model Eq. (1) in the absence of stimulation (arrowed black curve). Superimposing these two representations allows us to break down the different regions of the bifurcation diagram: when the black trajectory is in the region with low values of membrane potential *V*, i.e. the red part of the surface, then it is the interictal region. When the curve approaches the LP_1_ bifurcation point, then the trajectory jumps to the blue surface where the values of the membrane potential are high, which represents the ictal phase. Then the curve jumps back down at the HB_1_ bifurcation point. This abrupt jump is due to a periodic branch which emerges from HB_1_, forming a direct homoclinic connection to the S-shaped surface. For clarity, we have omitted the periodic branch in Fig. 5.B, which is further elaborated on in Supplemental Material S4. LP_1_ can be regarded as the potassium threshold for which the model produces a seizure if it is exceeded. The grey colored surface corresponds to the unstable states of the system that separate the basins of attraction of the ictal and interictal states.

Having established the different regions, in Fig. 5.C we study the solutions of the model subjected to LFES in the phase plane {[*K*^+^]_*o*_, [*Na*^+^]_*i*_}, indicating the threshold values obtained in Fig. 5.B (HB_1_, LP_1_, LP_2_) by horizontal lines for better readability. The half-planes below LP_2_ and above LP_1_ in Fig. 5.C belong to the interictal and the ictal region, respectively. There is a hysteresis effect due to the coexistence of the ictal and interictal regime between LP_1_ and LP_2_. If a trajectory approaches this interval from the interictal regime (below LP_2_), then it remains in the interictal regime; and conversely, if it approaches this interval from the ictal regime (above LP_1_), then it will remain in the ictal regime. In the top panels of Fig. 5.C, we plot several trajectories with different stimulation onsets (indicated by the dots of different colors). The blue cross indicates the critical stimulation onset established in Fig. 4. For any stimulation before this critical point, the LFES is effective, otherwise, it causes a premature seizure. For the case *A* = 10 mA, the stimulation pushes the system onto a smaller limit cycle (compared to the unperturbed system) that passes through the thresholds. Thus, for any stimulation below the critical stimulation amplitude, seizures are shorter with smaller amplitudes, and shorter ISIs. When *A* = 15 mA, for any stimulation arriving before the critical point of stimulation onset, the system stabilizes on a smaller limit cycle. This limit cycle is also observable in the case *A* = 30 mA. Since these limit cycles no longer pass through the LP_1_ point, they are no longer associated with seizure behavior, but instead, keep the system locked in a subpart of the interictal domain. These results corroborate our observations in Fig. 4, and further suggest that the stronger the stimulation (above *A* = 15 mA), the faster is the stabilization at this new stationary cycle.

In Fig. 4, we predicted that if we choose the right amplitude to induce a delay but stimulate too late, then we induce a single transient seizure before converging to the delay process. In Fig. 5.C we observe this behavior in the phase plane for *A* = 15 mA and *A* = 30 mA. The trajectory passes through the seizure domain once, before converging to the new attractor. This implies that the onset threshold relative to the last seizure only acts as a threshold for the initial conditions of the LFES. In fact, for any LFES above 15 mA, the delay appears with or without a single transient seizure.

The second and third row of panels in Fig. 5.C show the trajectories obtained from the same simulations as the first row, but with color coding of the current induced by the Na-K pump *I*_*pump*_, and the firing rate *υ*_*i*_, respectively. We can observe the activation and deactivation of the Na-K pump during a seizure: initially, the intracellular sodium concentration is low, and the pumps have very low activity. During the seizure, at the peak of activity, the pump reactivates due to the accumulated intracellular sodium, and decreases the extracellular potassium concentrations, which brings the neuronal population back to the baseline. When stimulating with *A* = 10 mA, this process also occurs during the appearance of the premature seizure. When the amplitude is large enough to delay seizures, the system converges onto a new attractor where the Na-K pump becomes increasingly active compared to the resting state without stimulation. The firing rate *υ*_*i*_ provides an indication of the effect of the stimulation protocol on the model. During the interictal phase, the firing rate is close to zero, whereas in the ictal phase it is close to the maximum *υ*_*max*_. When LFES is applied to the system, the firing rate shows a burst-like response to the stimulation pulses. However, the amplitude of these bursts only reaches its maximum when the stimulation is effective in delaying the subsequent seizure. The bursts of firing activity increase K and Na concentrations, which is counteracted by the Na-K pumps, thus establishing a new attractor state.

In summary, this analysis highlights the critical role of the Na-K pump in the processes induced by LFES, stabilizing ionic concentrations at relatively low values. We demonstrate that the observed phenomenon is not associated with epileptic activity. Instead, it represents an interictal dynamic that maintains the system at a safe distance from the seizure transition.

## Discussion

In this study, we aim to gain deeper insights into brain stimulation as a treatment for epilepsy, by employing a modified version of the Epileptor-2 model (25), calibrated with *in vitro* rat brain data. This work bridges the gap between experimental findings and theoretical insights, providing a deeper understanding of how LFES interacts with seizure dynamics at a mechanistic level. Leveraging experimental data from hippocampal rat slices under high-potassium condition, we reproduce key features of seizure-like events, such as their onset, duration, and ISIs, while also accounting for the variability observed across individual slices. The model not only recapitulates these experimental findings, but also offers novel insights into how LFES modulates ion dynamics, particularly through its interaction with the Na-K pump. This mechanism stabilizes ionic concentrations, thereby delaying seizure onset, maintaining the system in an interictal state. By systematically exploring the onset, amplitude, and duration of LFES, we identify optimal stimulation parameters maximizing its anti-seizure effects without inadvertently triggering epileptic activity.

The choice of applying LFES at 1 Hz was made to match the stimulation protocol used during data acquisition (19). This frequency has been widely adopted in previous studies as it has been shown to induce lasting changes in cortical excitability and to reduce seizure frequency in epilepsy models and clinical applications (40–42).

In the original Epileptor-2 model, the addition of LFES in the inter-seizure interval had either no effect, or caused premature seizures for any amplitude (see Supplemental Material S5). We therefore applied some modifications necessary to account for the seizure delay computationalcomputationaldynamics of the inhibitory neuronal population. The latter was considered to balance the dynamics of the excitatory population in the original model, which we replaced by an inhibitory population providing background inhibition at a constant firing rate (27). However, in Supplemental Material S6, we demonstrate that after removing inhibition completely from the modified Epileptor-2, the model still produces seizures, and LFES is effective in delaying them. This implies that the inhibitory population in our model is not crucial for the emergence of seizures or their delay.

There are some differences between the experimental protocol and the model setup. First, our ability to fit the model to the data is limited by the nature of the LFP, which is generated by multiple different processes (43, 44) that are difficult to disentangle, especially in terms of the amplitude of the signal. Therefore, while the data represent the LFP measured in the extracellular medium, the model describes the average membrane potential of the pyramidal cells. To obtain a match between the data and model, we aligned the temporal features of the model output with the experimentally observed seizure dynamics, such as seizure length and ISI, ignoring the amplitude of the signal. To further demonstrate the model’s validity, we reproduced experimental protocols in model simulations, such as the effect of step-wise increase of potassium concentration in the ACSF (Fig. 2B). Second, in the experiment the stimulation was performed on the Schaffer collaterals coming from the CA3 region extending towards CA1. This stimulation likely elicits spike volleys in the axons traveling to CA1, where they generate postsynaptic potentials at the excitatory population. The relationship between the amplitude of the electric field and the number of elicited spikes in the Schaffer collaterals (and the resulting postsynaptic potentials in CA1) is likely nonlinear. It is reasonable to assume that stimulation leads to a binary response at the level of single axons, where the electric field is either strong enough to elicit a spike, or too weak to do so. Because larger axons activate at smaller field strengths than smaller axons (45), the morphological heterogeneity of axons may result in a more graded response, at least in a narrow range of electric field strengths. Since the exact relationship between the electric field strength and the resulting PSPs in CA1 is unknown, we decided to directly use the amplitude of the postsynaptic current as model parameter.

We further examined optimal LFES in terms of stimulation onset, duration and amplitude. We identified critical values for onset and amplitude: if the onset occurs too late, or the amplitude is too low, then even a single stimulation pulse has an adverse effect and triggers a premature seizure. For onset times before the critical time point and amplitudes above the critical intensity, their exact values do not play a role in the delaying effect. However, the duration of the stimulation is a crucial parameter, with which the relative delay of the seizure increases linearly, which is corroborated by our experimental results. Thus, the LFES applied to the model led to the emergence of a new small periodic equilibrium during stimulation, if applied at the right time and amplitude. Given our discussion in the previous paragraph, in the experimental setup the critical stimulation amplitude is likely mediated by the structural properties of the Schaffer collaterals, and therefore doesn’t directly relate to the stimulation amplitude used in the model.

Finally, the mathematical model allowed us to analyse the processes underlying the emergence of seizures and the seizure delay due to LFES. Seizures in the Epileptor-2 model occur periodically, driven by intracellular sodium and extracellular potassium concentration dynamics. During the interictal period, intracellular sodium decreases and extracellular potassium increases. The latter happens due to diffusion of potassium from the potassium-enriched ACSF perfusing the slice. Due to the accumulation of extracellular potassium, the system eventually switches to the ictal state, which is characterized by high extracellular potassium, the depolarisation of the membrane, and an increase in the firing rate. The increased firing rate leads to an accumulation of intracellular sodium throughout the ictal phase, and a further accumulation of extracellular potassium in the first stage of the seizure. In the second stage of the seizure, when both sodium and potassium concentrations are sufficiently high, the sodium-potassium pumps activate and decreasing extracellular potassium (but barely affecting the sodium concentration, due to the volume ratio parameter), eventually lowering the membrane potential sufficiently for the system to reverse to the interictal phase. Thus, it appears that Na-K pumps are crucial for the termination of ictal phase because they lower the extracellular potassium concentration. During the interictal phase, however, the activity of the Na-K pumps is not enough to compensate for the potassium concentration increase due to the diffusion from the high-potassium ACSF. There is a time window during the interictal phase, when the system is in, or close to, the regime where sodium-potassium pumps are active. The stimulation protocol causes the neurons to fire and, thus, increases both sodium and potassium concentrations. The system then moves to a state in which the activity of the Na-K pump is counterbalanced by the stimulation, which is sufficient to suppress further increases in the extracellular potassium concentration and, thus, prevents spontaneous seizure occurrence.

The crucial role played by Na-K-pumps in seizure termination as well as for the ionic balance in general, is supported by experimental studies (46–51). Furthermore, inherited and acquired sodium or potassium-related channelopathies (52, 53) and genetic Na-K-pump dysfunctions can cause epilepsy (54). Anti-seizure medications aimed at restoring normal Na-K-ATPase function might alleviate symptoms in patients with pump-related mutations (47, 52). From this perspective, Na-K-pumps emerge as promising strategic targets for therapeutic interventions aimed at modulating seizure activity. For instance, by enhancing or restoring Na-K-ATPase function, it might be possible to mitigate hyperexcitability and promote seizure termination, potentially offering novel avenues for treatment, especially in cases where pump dysfunction contributes to the disease. These hypotheses can first be experimentally tested in vitro, by selectively increasing or decreasing Na-K-pump activity and assessing the resulting effects on seizure dynamics. Such approaches have already been employed, where Na-K-pump activity was experimentally manipulated to investigate its impact on seizure dynamics. For example, blocking Na-K-pumps with ouabain elicits long-lasting seizures in otherwise healthy tissue (55). Such experimental strategies provide a valuable framework to validate our model’s predictions regarding the contribution of Na-K-pump dynamics to seizure generation and termination.

In our computational study, the results were obtained using a model that reproduces in vitro-like conditions, specifically based on data and observations from animal brain slice preparations. While the model itself is grounded in animal data, the fundamental biophysical mechanisms it captures — in particular, the role of Na-K-pumps in maintaining ionic homeostasis and contributing to seizure termination — are highly conserved across mammalian species, including humans (46, 51, 56, 57). This suggests that Na-K-pump-related processes could represent a common pathogenic factor underlying epileptic activity in both experimental models and human epilepsy.

The study from which the experimental data are taken (19), investigates the LFES effect by using a phenomenological modeling approach based on a reduced version of the phenomenological Epileptor model (58). This line of work was further extended employing a theoretical framework rooted in phase response analysis of planar relaxation oscillators (20). These models offer mathematical insight into how the slow dynamics of the system shape seizure generation and its modulation by external perturbations, such as LFES. However, being purely phenomenological, they do not explicitly incorporate biophysical variables that could explain the underlying mechanisms of stimulation at the cellular or network level. Other modeling studies have also investigated how LFES modulates epilepsy, such as the role of presynaptic GABA_*B*_ receptors within a phenomenological framework (12), or how thalamic stimulation at different frequencies can modulate neocortical activity (24), shedding light on the frequency-dependent nature of stimulation effects. Our work is therefore novel in that it explicitly addresses the question of which minimal biophysical mechanisms are necessary for LFES to modulate seizure activity. To this end, we develop and analyze a minimal biophysical model based on neural mass dynamics. This modeling strategy aims to bridge the gap between purely phenomenological descriptions and detailed conductance-based models, providing a mechanistic understanding of LFES within a simplified yet biophysically grounded framework. By identifying the essential processes required for LFES efficacy, our approach contributes to clarifying the causal mechanisms of stimulation and offers a theoretical foundation for optimizing stimulation protocols in epilepsy.

## Conclusion

In conclusion, this work provides a significant advancement in understanding the effects of LFES on seizure dynamics by modifying the Epileptor-2 model to better align with experimental data. By integrating parameters from in vitro hippocampal recordings, the model uncovers a key mechanism through which LFES modulates the Na-K pump, stabilizing ionic concentrations and delaying seizure onset. The identification of critical amplitude and duration parameters for LFES offers new insights into optimizing this therapeutic approach. These findings bridge the gap between theoretical modeling and experimentation, providing a robust framework for refining stimulation strategies in epilepsy treatment.

## ACKNOWLEDGEMENTS

This work was supported by ERDF-Project Brain dynamics, No. CZ.02.01.01/00/22_008/0004643, by project nr. LX22NPO5107 (MEYS): Financed by EU – Next Generation EU, Czech Science Foundation (21-17564S), a Lumina-Quaeruntur fellowship (LQ100302301) by the Czech Academy of Sciences (awarded to HS), and the long-term strategic development financing of the Institute of Computer Science (RVO:67985807) of the Czech Academy of Sciences, and Charles University grant PRIMUS/23/MED/011.

We would like to thank Thomas R. Knösche for valuable discussions.

## Supplemental Material 1: Data Analysis - Inter-Seizure-Intervals (ISI)

**Figure S1:**
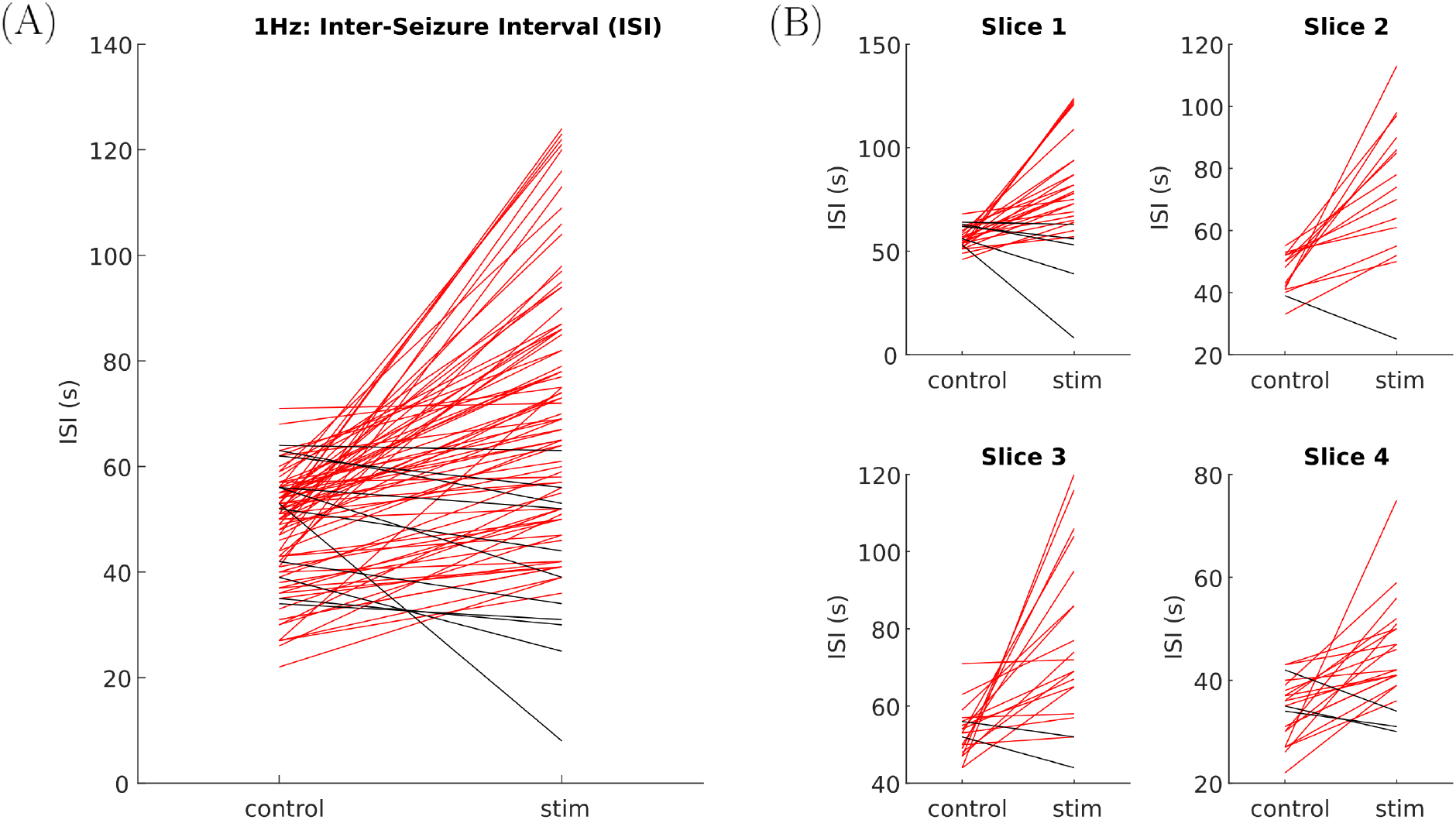
(A) Inter-seizure-interval across slices. Prolonged ISIs under stimulation are marked in red, shortened ISIs in black. Most ISIs are prolonged (i.e. seizure delay). (B) Inter-seizure-interval per slice. All slices show the delay effect. Note the variation in slice-specific ISI. The seizure delay (marked in red), as induced by 1 Hz stimulation, is clearly visible on the group level, as well as in the individual slices.

## Supplemental Material 2: LFES efficiency regions in Modified Epileptor-2

**Figure S2:**
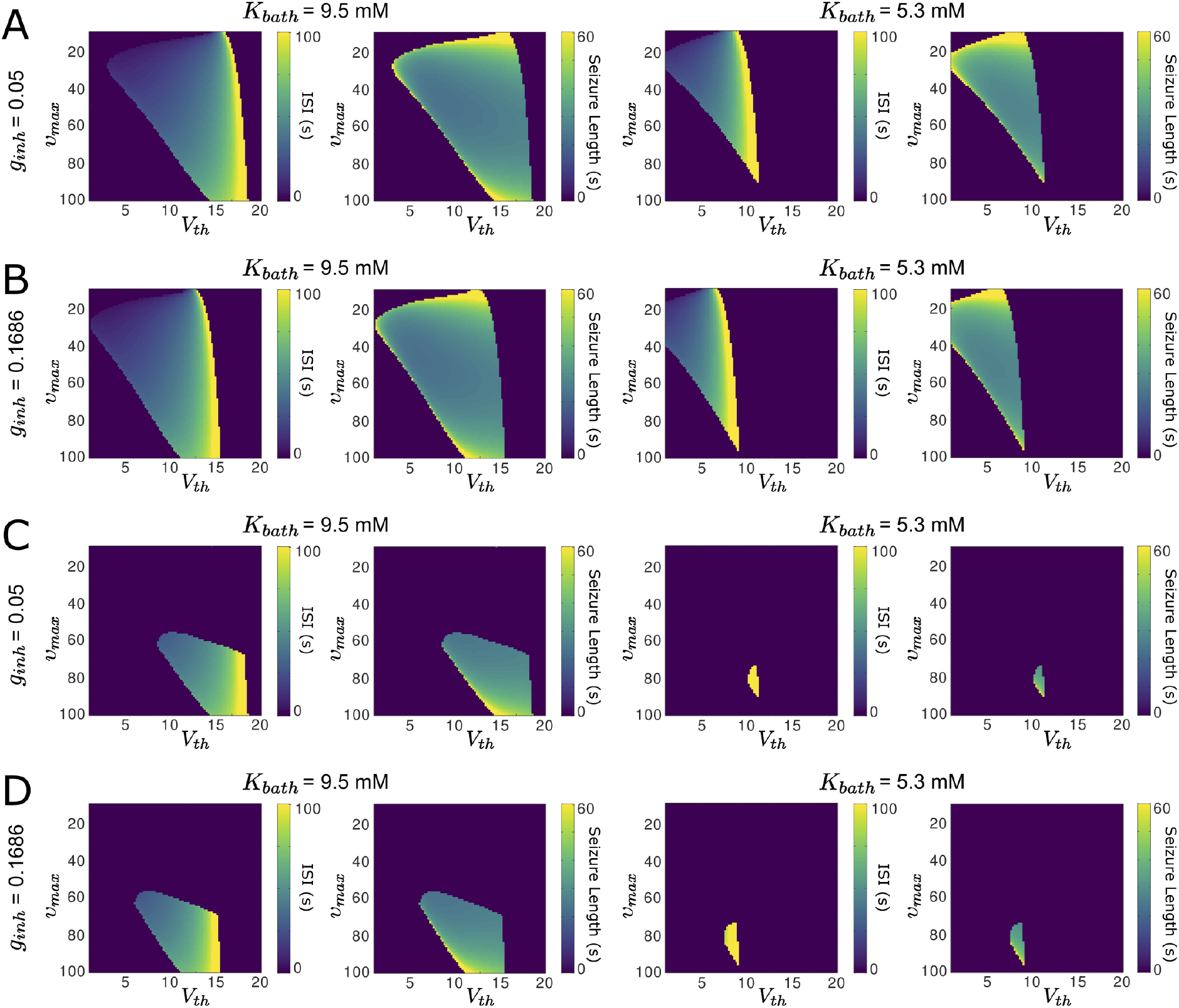
Heatmaps of ISI and seizure duration with respect to the parameters *V*_*th*_, *υ*_*max*_ and *g*_*inh*_, for slice 1 (*K*_*bath*_ = 9.5) and 4 (*K*_*bath*_ = 5.3). (A-B) Heatmaps for *g*_*inh*_ = 0.1686 and *g*_*inh*_ = 0.05 respectively. (C-D) Same activity maps, however, the pixels for which the model exhibits seizures, but where the LFES is not effective have been set to 0. LFES fails to delay seizures in slice 4 (see Fig 3), prompting an investigation using activity maps to assess the effects of inhibitory current and key parameters on seizure dynamics. These maps reveal how inhibition influences seizure occurrence, duration, and spacing under slice 1 and 4 conditions. Increasing inhibition shifts the seizure-prone region to higher threshold values, enabling better data fitting for slice 4 with *g*_*inh*_ = 0.1686. However, while slice 1 allows both accurate data fitting and effective LFES, no parameter set achieves this for slice 4, highlighting limitations in the model’s ability to reproduce both seizure dynamics and stimulation effects. For the specific parameters for each slice, see the main result Table 3.

## Supplemental Material 3: Data Analysis - Stimulation parameters

**Figure S3:**
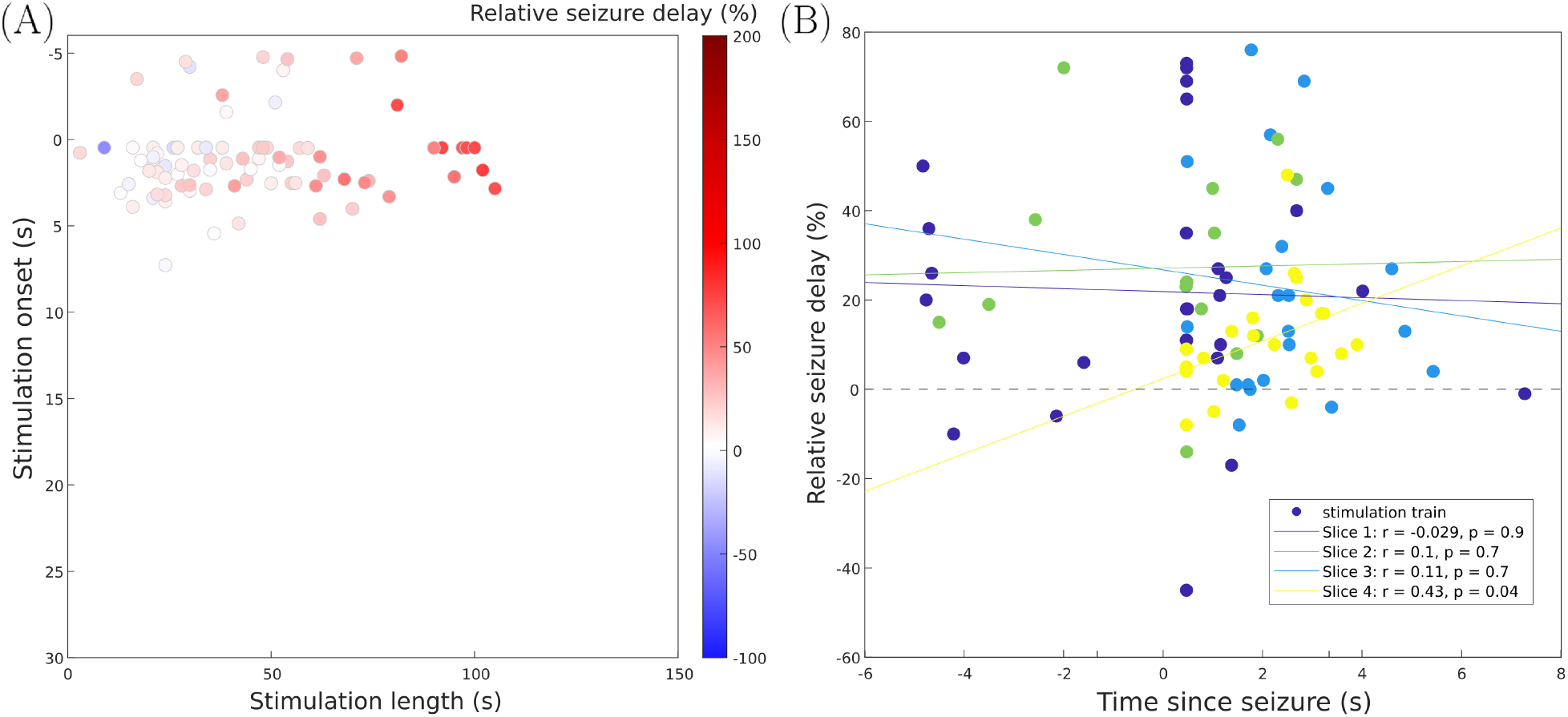
(A) Stimulation onset time expressed as time since last seizure and stimulation length, displayed in accordance with Fig. 4.C of the main manuscript. Most of the parameter space explored in Fig. 4.C was not covered in the experimental dataset. Stimulation onset in the dataset was centered around last seizure offset. For stimulation lenth, the experimental observation matched the model prediction: longer stimulation time led to longer ISI as compared to the control ISI. (B) Relationship of stimulation parameters and seizure delay in each of the slices (color-coded). For the narrow parameter range explored, stimulation onset time did not seem to correlate with with the seizure delaying effect of LFES. Due to data limitations, a thorough exploration of the parameter space, as for the model in Fig. 4 of the main manuscript, was not possible. Stimulation amplitude was not available. Stimulation onset time varied only in a small range close to previous seizure end in panel (A). In this time window we did not see a significant correlation between seizure delay and stimulation onset time (“Time since seizure”) in panel (B).

## Supplemental Material 4: {[*K*^+^]_*o*_, *V*}-bifurcation diagram

**Figure S4:**
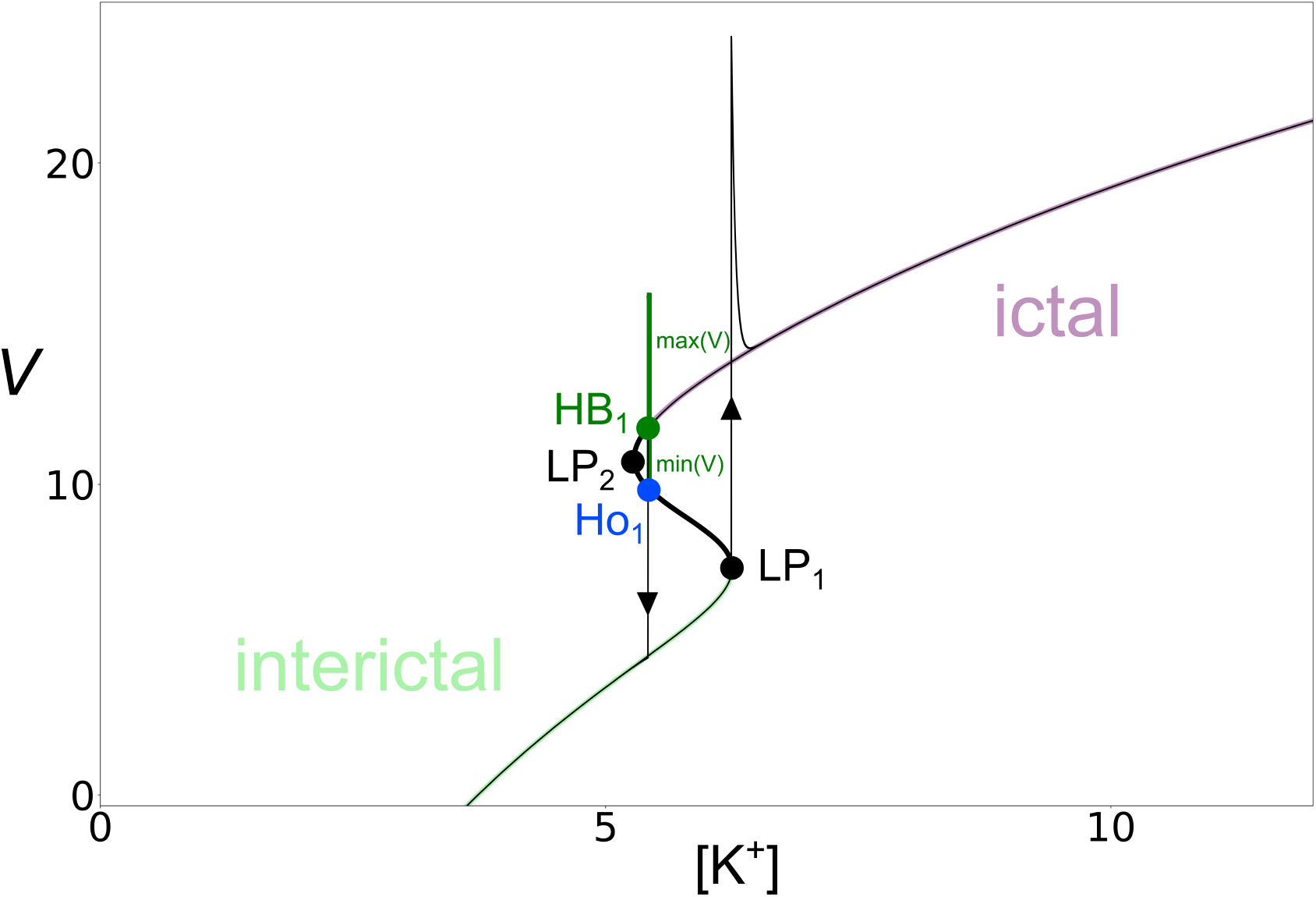
[*K*^+^]_*o*_, *V* bifurcation diagram obtained from the fast-subsystem (Eq. Eq. (7)), overlaid with a non-stimulated solution obtained from Eq. Eq. (1) (black arrowed curve). For the superposition to be in perfect adequacy, we impose *ε* = 0.003 parameter in front of the equation of [*K*^+^], in such a way, they are slower and follow the bifurcation structure. The stable states are the light green and purple curves, which refer respectively to the inter-ictal region and the ictal region. The black curve represents the unstable states. HB denotes Hopf bifurcation, LP, fold bifurcation, and Ho, homoclinc bifurcation. The deep green curve refers to the periodic branch. This figure highlights the periodic branch arising from the Hopf point, and the presence of a homoclinic bifurcation point when it is interrupted in the unstable branch of the S-shaped stationary point curve. The verticality and explosiveness of the periodic branch therefore do not allow the existence of a periodic solution, even with slow trajectories. Thus, the solution that crosses the Hopf point after passing the ictal phase, jumps directly to the interictal phase, passing through the point Ho_1_.

## Supplemental Material 5: Original Epileptor-2 model study

**Table S1:**
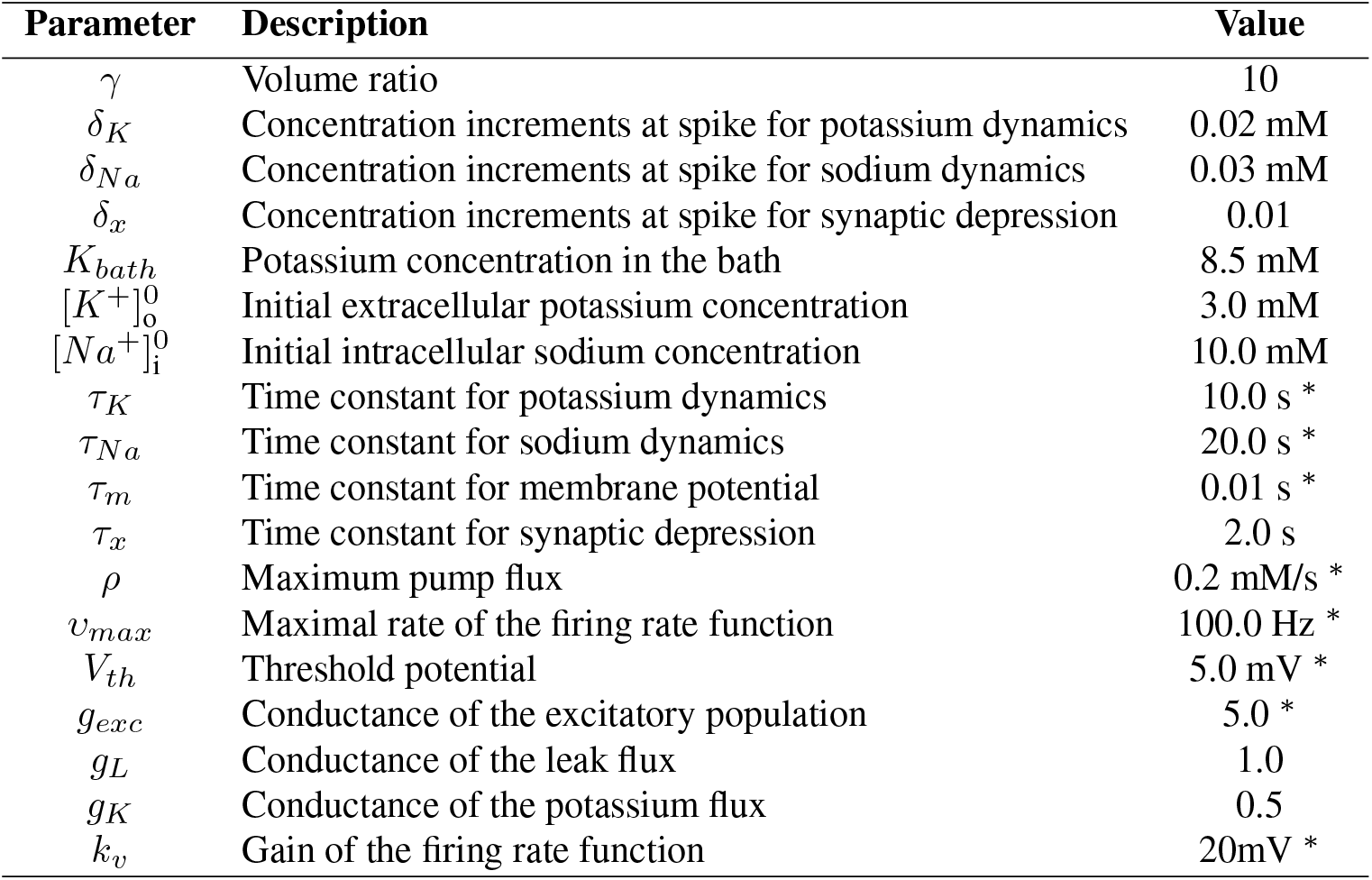
Original parameter set for the Epileptor-2 model. The parameters marked with ^∗^ are different from modified Epileptor-2. The original Epileptor-2 equations are defined in (25). The decision to modify certain parameters was driven by the aim to obtain a noise-free model that reliably generates periodic seizures. Additionally, we sought to ensure that the membrane potential amplitude of the model closely matches the experimental data.

**Figure S5:**
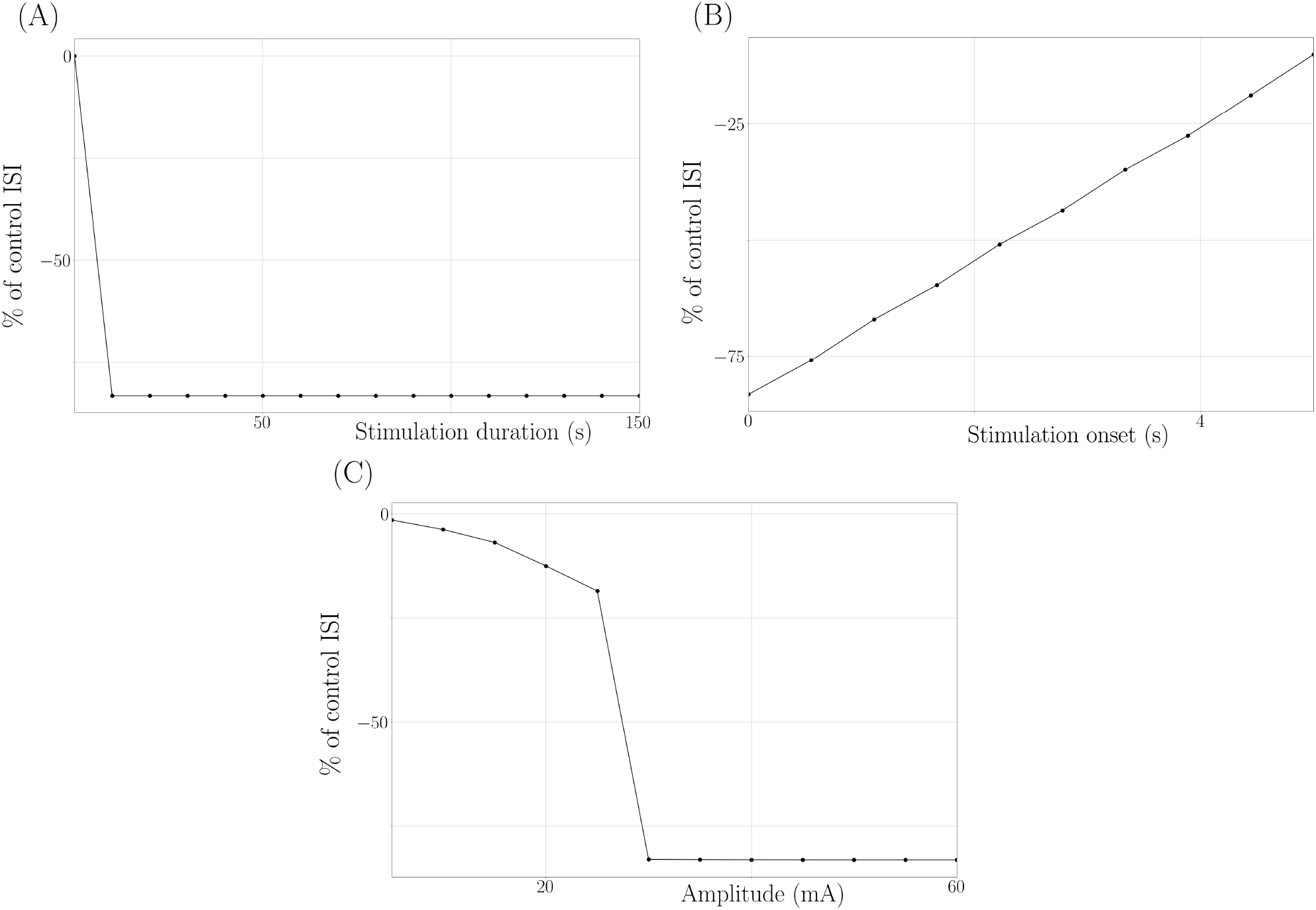
LFES effects on the Epileptor-2 model. Stimulated ISI expressed as a percentage relative to the unstimulated ISI, with respect to (A) stimulation duration (s), (B) stimulation shift (s), and (C) amplitude (mA). Results show that stimulation generally shortens the ISI, with earlier stimulations leading to earlier subsequent seizures, and that increasing amplitude beyond 30mA does not further delay seizure onset. The parameter set is described in Table S1.

## Supplemental Material 6: Removing the inhibition dependency

**Figure S6:**
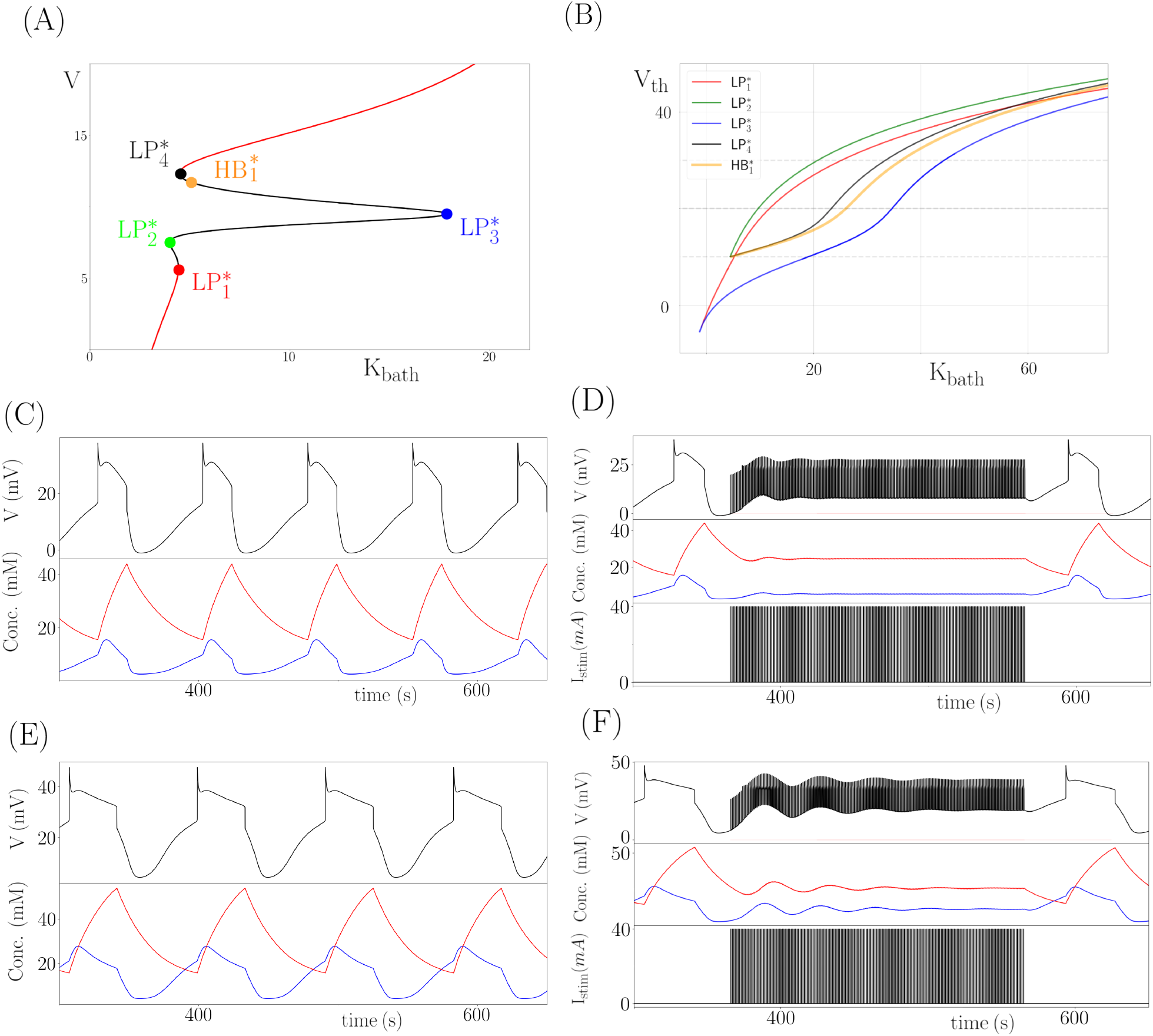
(A) Bifurcation diagram of *V* with respect to *K*_*bath*_ of the system Eq. Eq. (1), but with *g*_*inh*_ = 0. Black (resp. red) segments of the *S*-shaped curve of equilibria denote unstable (resp. stable) families of equilibria. (B) {*K*_*bath*_, *V*_*th*_}-parameter bifurcation diagram. HB^∗^ denotes Hopf bifurcation and LP^∗^, fold bifurcation point. The three horizontal grey dashed lines are the different scenario we are studying in this figure : *V*_*th*_ = 10, 20, 30 mV. We also show the potential and ion dynamic time series for *V*_*th*_ = 20 mV and *K*_*bath*_ = 15mM ((C) unstimulated, (D) stimulated case); and for *V*_*th*_ = 30 mV and *K*_*bath*_ = 29mM ((E) unstimulated, (F) stimulated case). The potassium ion dynamic is the blue curve, and the sodium one is the red curve. The bifurcation diagram (A) shows that LP1 and LP4 are too close to allow clear seizure expression, prompting further analysis in the {*K*_*bath*_, *V*_*th*_} space. By selecting *V th* = 20*mV* and *V th* = 30*mV*, we demonstrate that seizure delay occurs without inhibition, justifying the inclusion of an inhibitory term to better reflect neuronal interactions.

